## supplementary figures and tables for "Chemical, physical and biological triggers of evolutionary conserved Bcl-xL-mediated apoptosis"

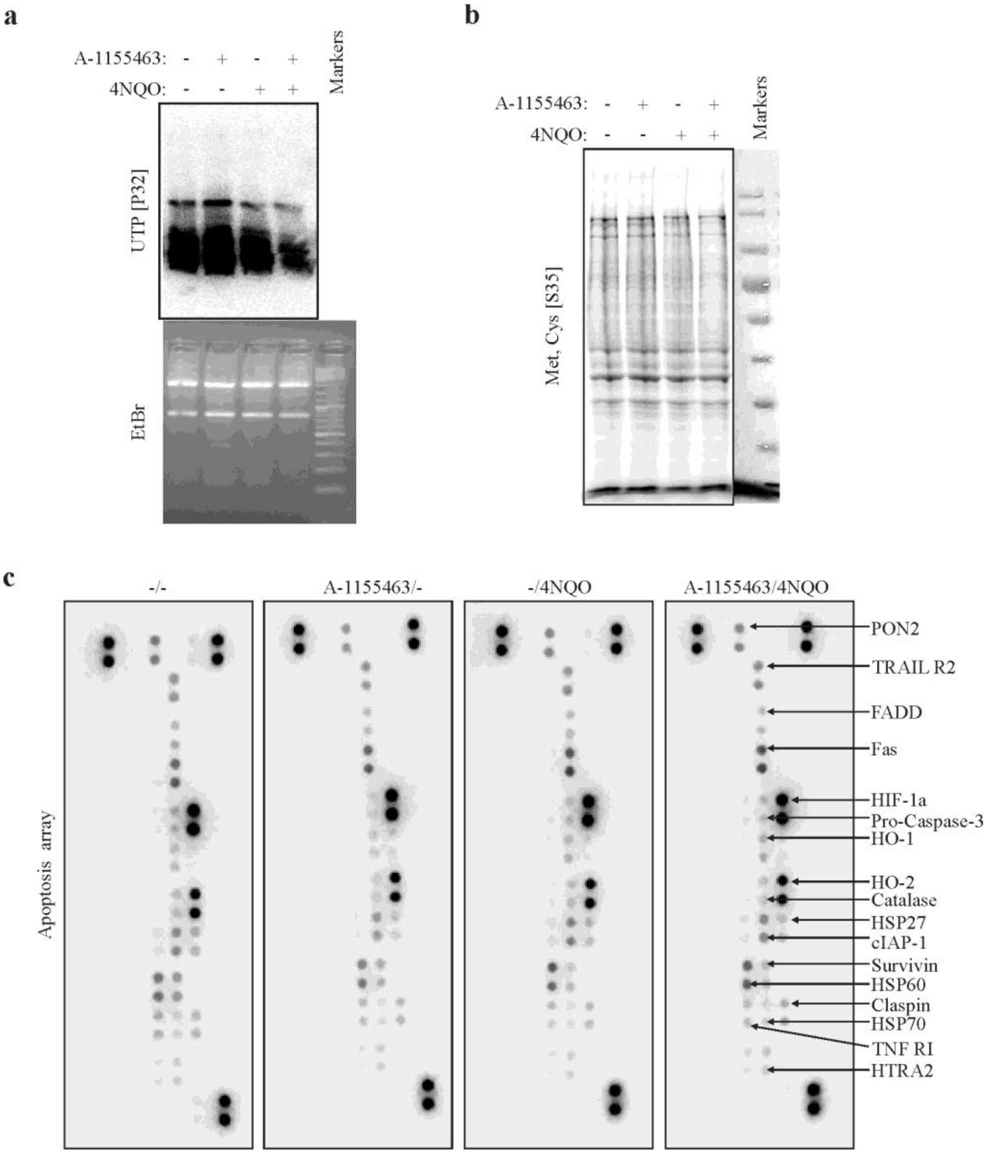


Figure S1. **Effect of 4NQO, A-1155463 or their combination on general transcription, translation, and specific apoptotic proteins**. (**a**) RPE cells were treated with 1 μM 4NQO, 1 μM A-1155463 or their combination. Control cells were treated with 0.1% DMSO. [alpha-P32]UTP was added to cell culture medium to label newly transcribed cellular RNA. Total RNA was isolated and subjected to agarose gel electrophoresis followed by radioautography. (**b**) RPE cells were treated as for (a). [$^{35}$S] methionine and cysteine were added to methionine- and cysteine-free culture medium to label newly synthetized cellular proteins. Cells were lysed and proteins were separated on SDS-polyacrylamide gel. $^{35}$S-labelled proteins were detected using radioautography. (**c**) RPE cells were treated as for (a). Relative levels of apoptosis-related proteins were determined using proteome profiler human apoptosis kit (n=2).

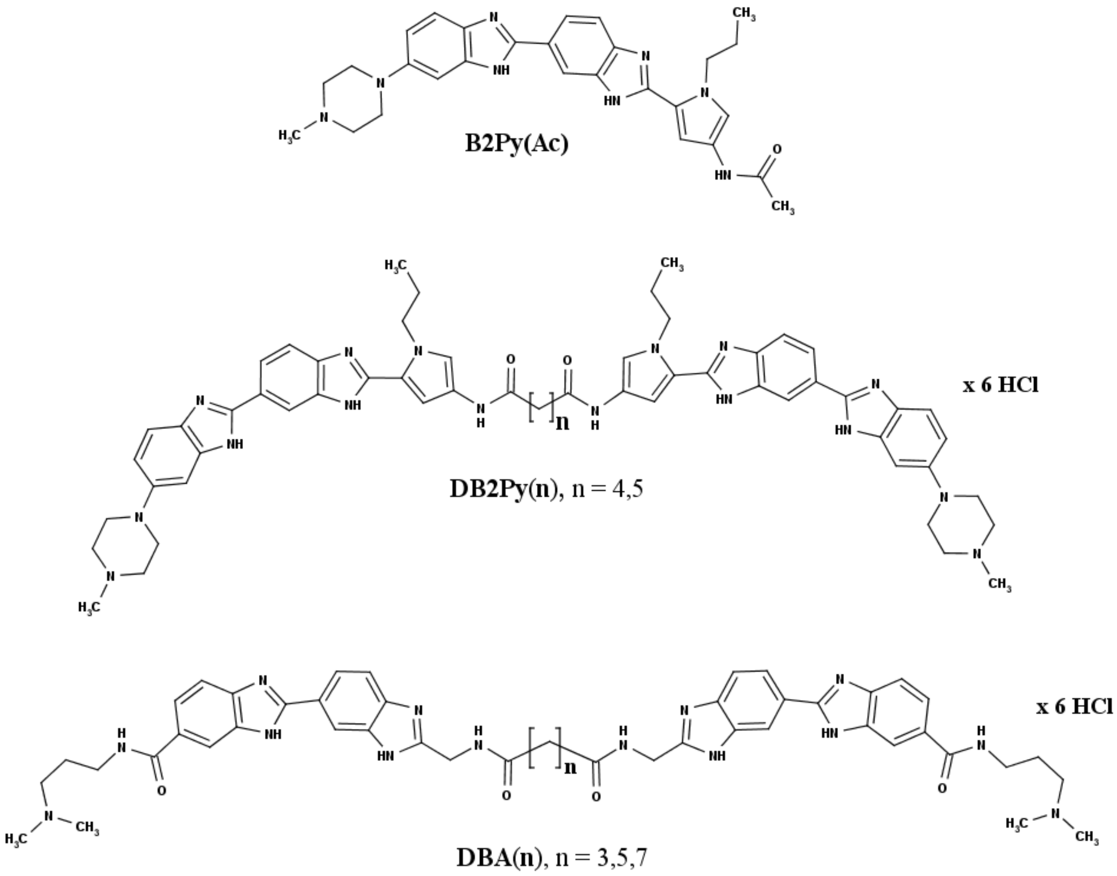


Figure S2. **Chemical structures of monomeric MB2Py(Ac) and dimeric DB2Py(n) bisbenzimidazole-pyrroles, as well as dimeric bisbenzimidazoles DBA(n).**

Table S1. **The active compounds of 48 commonly prescribed drugs in Norway, their suppliers and catalogue numbers.**

| Drug | CAS | MW | Formula | Cat N | Purity, % | Supplier |
|---|---|---|---|---|---|---|
| 17α-Ethynylestradiol | 57-63-6 | 296 | C20H24O2 | E4876-100MG | ≥98 | Sigma Aldrich |
| 4-Acetamidophenol | 103-90-2 | 151 | C8H9NO2 | 102330050 | 98 | Acros Organics |
| Acetylsalicylic acid | 50-78-2 | 180 | C9H8O4 | AC158180500 | 99 | Acros Organics |
| Amlodipine | 88150-42-9 | 409 | C26H31ClN2O8S | CAYM14838 | ≥98 | Cayman Chemicals |
| Atorvastatin | 134523-03-8 | 559 | C33H35FN2O5 | CAYM10493 | ≥98 | Cayman Chemicals |
| Bumetanide | 28395-03-1 | 364 | C17H20N2O5S | CAYM14630 | ≥98 | Cayman Chemicals |
| Candesartan | 139481-59-7 | 440 | C24H20N6O3 | sc-217825 | ≥98 | Santa Cruz Biotechnology |
| Cetirizin | 83881-52-1 | 389 | C21H27Cl3N2O3 | 89126-50MG | ≥98 | Sigma Aldrich |
| Cyanocobalamin | 68-19-9 | 1355 | C63H88CoN14O14P | DRE-C11798500 | | LGC Standards |
| Desloratadine | 100643-71-8 | 311 | C19H19ClN2 | CAYM16931 | ≥98 | Cayman Chemicals |
| Desogestrel | 54024-22-5 | 310 | C22H30O | CAYM23651 | ≥95 | Cayman Chemicals |
| D-Pantothenic acid | 79-83-4 | 219 | C9H17NO5 | HY-B0430 | ≥98 | MedChemExpress |
| Drospirenone | 67392-87-104 | 367 | C24H30O3 | CAYM23347 | ≥98 | Cayman Chemicals |
| Enalapril | 75847-73-3 | 376 | C20H28N2O5 | J60750.03 | ≥97 | Alfa Aesar |
| Escitalopram | 128196-01-0 | 324 | C20H21FN2O | CAYM22405 | ≥98 | Cayman Chemicals |
| Esomeprazole | 161973-10-0 | 767 | C34H42MgN6O9S2 | CAYM17326 | ≥95 | Cayman Chemicals |
| Etonogestrel | 54048-10-1 | 324 | C22H28O2 | CAYM21062 | ≥98 | Cayman Chemicals |
| Fluticasone propionate | 80474-14-2 | 445 | C25H31F3O5S | 462101000 | ≥96 | Acros Organics |
| Folic acid | 59-30-3 | 441 | C19H19N7O6 | J62937.06 | ≥97 | Alfa Aesar |
| Furosemide | 54-31-9 | 331 | C12H10ClN2O5S | 448970010 | ≥97 | Acros Organics |
| Hydroxocobalamin | 13422-5 51-0 | 1346 | C62H89CoN13O15P | CAYM24099 | ≥95 | Cayman Chemicals |
| Insulin aspart | 116094-23-6 | 5826 | C256H387N65O79S6 | EPY0000349 | | LGC Standards |
| Lercanidipine | 132866-11-6 | 612 | C36H41N3O6 | HY-B0612A | 98.5 | MedChemExpress |
| Levonorgestrel | 797-63-7 | 312 | C21H28O2 | CAYM10006 | ≥95 | Cayman Chemicals |
| Levothyroxine | 25416-653 | 817 | C15H12I4NNaO5 | FT48192 | ≥97 | Carbosynth |
| Losartan | 114798-26-4 | 423 | C22H23ClN6O | FL39656 | ≥97 | Carbosynth |
| Metformin | 1115-70-4 | 166 | C4H12ClN5 | sc-202000 | ≥99 | Santa Cruz Biotechnology |
| Metoprolol | 51384-51-1 | 267 | C15H25NO3 | sc-264643 | 97 | Santa Cruz Biotechnology |
| Mometasone furoate | 83919-23-7 | 521 | C27H30Cl2O6 | CAYM21365 | ≥98 | Cayman Chemicals |
| Naproxen | 22204-53-1 | 230 | C14H14O3 | CAYM70290 | ≥99 | Cayman Chemicals |
| Nicotinic acid | 59-67-6 | 123 | C6H5NO2/HOOC5H4N | 128290050 | 99.5 | Acros Organics |

| | | | | | | |
|---|---|---|---|---|---|---|
| Nifedipine | 21829-25-4 | 346 | C17H18N2O6 | CAYM11106 | ≥98 | Cayman Chemicals |
| Pantoprazole | 102625-70-7 | 383 | C16H15F2N3O4S | CAYM21345 | ≥98 | Cayman Chemicals |
| Prednisolone | 50-24-8 | 360 | P6004 | P6004 | ≥98 | SigmaAldrich |
| Pyridoxine | 58-56-0 | 206 | C8H12ClNO3 | A12041.14 | ≥98 | WVR |
| Ramipril | 87333-19-5 | 417 | C23H32N2O5 | FC27676 | ≥98 | Cymit Quimica |
| Riboflavin | 83-88-5 | 376 | C17H20N4NaO9P | A11764.14 | 98 | Alfa Aesar |
| Salbutamol | 18559-94-9 | 239 | C13H21NO3 | CAYM21003 | ≥98 | Cayman Chemicals |
| Salmeterol | 89365-50-4 | 416 | C25H37NO4 | HY-14302 | 99.7 | MedChemExpress |
| Sertraline | 79559-97-0 | 306 | C17H18Cl3N | 462190010 | ≥98 | Acros Organics |
| Simvastatin | 79902-63-9 | 419 | C25H38O5 | 458840010 | 98 | Acros Organics |
| Tamsulosin | 106463-17-6 | 445 | C20H29ClN2O5S | CAYM24020 | ≥98 | Cayman Chemicals |
| Thiamine | 67-03-8 | 337 | HC12H17ON4SCl2 | 148990100 | 99 | Acros Organics |
| Valsartan | 137862-53-4 | 436 | C24H29N5O3 | sc-220362 | ≥98 | Santa Cruz Biotechnology |
| Venlafaxine | 99300-78-4 | 277 | C17H27NO2 | HY-B0196A | 98 | MedChemExpress |
| Vitamin D2 | 50-14-6 | 397 | C28H44O | CAYM11791 | ≥98 | Cayman Chemicals |
| Vitamin D3 | 67-97-0 | 385 | C27H44O | CAYM11792 | ≥98 | Cayman Chemicals |

Table S2**. The synergy scores of combinations of A-1331852 with 527 anticancer agents**.

| Drug.combination | Synergy.score | Most.synergistic.area.score | Method |
|---|---|---|---|
| Mivebresib | 20,199 | 29,23 | ZIP |
| Cabazitaxel | 16,342 | 29,985 | ZIP |
| Indibulin | 15,39 | 30,956 | ZIP |
| Pictilisib | 15,269 | 24,627 | ZIP |
| GSK-461364 | 14,805 | 31,028 | ZIP |
| SN-38 | 14,752 | 21,513 | ZIP |
| Altiratinib | 14,324 | 24,411 | ZIP |
| Cisplatin | 13,888 | 25,177 | ZIP |
| Amsacrine | 13,837 | 22,067 | ZIP |
| Vinorelbine | 12,612 | 22,828 | ZIP |
| dBET1 | 12,067 | 24,377 | ZIP |
| Eltanexor | 11,872 | 18,343 | ZIP |
| RGFP966 | 11,044 | 14,619 | ZIP |
| Tamatinib | 10,915 | 21,034 | ZIP |
| Etoposide | 10,461 | 13,868 | ZIP |
| Cerdulatinib | 9,968 | 21,125 | ZIP |
| AMG-232 | 9,93 | 17,457 | ZIP |
| CC-115 | 9,877 | 14,517 | ZIP |
| NVP-LCL161 | 9,874 | 16,346 | ZIP |
| GSK269962 | 9,864 | 20,575 | ZIP |
| Mitoxantrone | 9,676 | 14,308 | ZIP |

| | | | |
|---|---|---|---|
| S-63845 | 9,488 | 29,158 | ZIP |
| Dactinomycin | 9,241 | 14,351 | ZIP |
| Dinaciclib | 9,235 | 14,165 | ZIP |
| THZ2 | 9,073 | 19,607 | ZIP |
| Docetaxel | 9,07 | 13,123 | ZIP |
| NVP-BHG712 | 8,715 | 17,165 | ZIP |
| Resminostat | 8,691 | 15,241 | ZIP |
| Foretinib | 8,633 | 13,023 | ZIP |
| Alvocidib | 8,593 | 20,266 | ZIP |
| Idasanutlin | 8,578 | 17,651 | ZIP |
| Eribulin | 8,482 | 17,324 | ZIP |
| CPI-360 | 8,414 | 12,401 | ZIP |
| BAY-1436032 | 8,218 | 11,584 | ZIP |
| GSK2801 | 8,058 | 13,099 | ZIP |
| UCN-01 | 8,037 | 14,593 | ZIP |
| AZD-8186 | 8,015 | 11,721 | ZIP |
| SGC0946 | 7,982 | 12,598 | ZIP |
| NVP-CGM097 | 7,869 | 13,193 | ZIP |
| BGB324 | 7,79 | 8,319 | ZIP |
| Tipifarnib | 7,762 | 13,709 | ZIP |
| Triciribine | 7,758 | 15,978 | ZIP |
| Birinapant | 7,748 | 12,656 | ZIP |
| Tofacitinib | 7,744 | 11,688 | ZIP |
| Sitravatinib | 7,681 | 10,744 | ZIP |
| ABC294640 | 7,679 | 11,178 | ZIP |
| Fludarabine | 7,585 | 11,915 | ZIP |
| GSK2656157 | 7,527 | 11,931 | ZIP |
| Rigosertib | 7,522 | 13,082 | ZIP |
| Tepotinib | 7,491 | 12,1 | ZIP |
| Alpelisib | 7,445 | 13,287 | ZIP |
| Serabelisib | 7,427 | 14,818 | ZIP |
| SB 743921 | 7,391 | 14,216 | ZIP |
| Pravastatin | 7,371 | 12,415 | ZIP |
| TRAM-34 | 7,344 | 11,329 | ZIP |
| PHA 408 | 7,323 | 13,274 | ZIP |
| CUDC-907 | 7,3 | 12,747 | ZIP |
| Tosedostat | 7,294 | 11,356 | ZIP |
| Selinexor | 7,29 | 13,122 | ZIP |
| BX-912 | 7,269 | 13,835 | ZIP |
| Sirolimus | 7,262 | 11,243 | ZIP |
| GSK923295 | 7,257 | 13,474 | ZIP |
| GSK650394 | 7,149 | 13,518 | ZIP |

| | | | |
|---|---|---|---|
| Abexinostat | 7,11 | 12,871 | ZIP |
| SAR405838 | 7,083 | 11,849 | ZIP |
| OTS-964 | 6,985 | 11,774 | ZIP |
| Pixantrone | 6,985 | 9,714 | ZIP |
| Tucatinib | 6,955 | 12,465 | ZIP |
| BMS-777607 | 6,955 | 11,733 | ZIP |
| Neratinib | 6,949 | 12,378 | ZIP |
| PFI-1 | 6,858 | 12,223 | ZIP |
| Ridaforolimus | 6,812 | 11,926 | ZIP |
| Deferoxamine | 6,795 | 10,842 | ZIP |
| PTC-209 | 6,722 | 9,169 | ZIP |
| SGC-CBP30 | 6,713 | 12,109 | ZIP |
| SNS-032 | 6,703 | 16,743 | ZIP |
| GSK2636771 | 6,703 | 12,068 | ZIP |
| AZD-5438 | 6,687 | 23,811 | ZIP |
| BMS-754807 | 6,68 | 10,621 | ZIP |
| ABT-751 | 6,614 | 12,751 | ZIP |
| GDC-0084 | 6,591 | 11,626 | ZIP |
| CUDC-305 | 6,59 | 16,263 | ZIP |
| Vincristine | 6,481 | 17,258 | ZIP |
| Pemetrexed | 6,437 | 10,575 | ZIP |
| TAK-901 | 6,402 | 11,856 | ZIP |
| Filanesib | 6,359 | 11,774 | ZIP |
| Prexasertib | 6,357 | 8,931 | ZIP |
| AZD1480 | 6,35 | 10,088 | ZIP |
| Itraconazole | 6,343 | 13,509 | ZIP |
| CPI-613 | 6,342 | 12,839 | ZIP |
| AZD0156 | 6,3 | 11,07 | ZIP |
| Paclitaxel | 6,201 | 11,743 | ZIP |
| Bleomycin | 6,187 | 10,261 | ZIP |
| Silmitasertib | 6,168 | 10,157 | ZIP |
| PIM-447 | 6,165 | 9,489 | ZIP |
| FRAX486 | 6,164 | 10,067 | ZIP |
| PF-06463922 | 6,159 | 11,485 | ZIP |
| PF-04708671 | 6,131 | 12,511 | ZIP |
| Talmapimod | 6,084 | 11,244 | ZIP |
| PF-00477736 | 6,074 | 10,457 | ZIP |
| Duvelisib | 6,038 | 9,762 | ZIP |
| CPI-0610 | 6,023 | 11,089 | ZIP |
| AZD8055 | 6,02 | 8,489 | ZIP |
| Birabresib | 6,018 | 8,285 | ZIP |
| Tubacin | 5,993 | 11,07 | ZIP |

| | | | |
|---|---|---|---|
| PF-00562271 | 5,989 | 11,042 | ZIP |
| BCI | 5,96 | 10,515 | ZIP |
| Baricitinib | 5,931 | 10,093 | ZIP |
| Saridegib | 5,914 | 11,29 | ZIP |
| Resiquimod | 5,909 | 10,352 | ZIP |
| GSK-690693 | 5,896 | 13,948 | ZIP |
| PF-03758309 | 5,891 | 8,958 | ZIP |
| Vidofludimus | 5,888 | 10,344 | ZIP |
| Luminespib | 5,884 | 12,23 | ZIP |
| Doxorubicin | 5,826 | 10,11 | ZIP |
| Lucitanib | 5,802 | 11,536 | ZIP |
| C646 | 5,802 | 10,622 | ZIP |
| Litronesib | 5,798 | 13,728 | ZIP |
| 4-hydroxytamoxifen | 5,778 | 10,213 | ZIP |
| Afuresertib | 5,758 | 8,562 | ZIP |
| Temsirolimus | 5,757 | 8,186 | ZIP |
| AT13148 | 5,732 | 9,123 | ZIP |
| Icotinib | 5,729 | 9,575 | ZIP |
| Tivozanib | 5,725 | 7,306 | ZIP |
| Taselisib | 5,714 | 8,788 | ZIP |
| GSK343 | 5,702 | 9,563 | ZIP |
| NVP-BGT226 | 5,677 | 10,741 | ZIP |
| Abemaciclib | 5,67 | 9,136 | ZIP |
| ENMD-2076 | 5,662 | 11,569 | ZIP |
| Omacetaxine | 5,587 | 7,88 | ZIP |
| Veliparib | 5,583 | 11,077 | ZIP |
| Fostamatinib | 5,556 | 7,854 | ZIP |
| Milciclib | 5,535 | 10,821 | ZIP |
| Tanzisertib | 5,51 | 10,642 | ZIP |
| A-419259 | 5,441 | 10,957 | ZIP |
| Ripasudil | 5,372 | 13,624 | ZIP |
| Gilteritinib | 5,372 | 8,304 | ZIP |
| GSK-1070916 | 5,37 | 8,825 | ZIP |
| Molibresib | 5,354 | 10,706 | ZIP |
| Ivosidenib | 5,309 | 11,378 | ZIP |
| Rabusertib | 5,257 | 7,699 | ZIP |
| NMS-873 | 5,251 | 12,94 | ZIP |
| Omipalisib | 5,232 | 8,122 | ZIP |
| Niraparib | 5,177 | 10,037 | ZIP |
| ARV-825 | 5,164 | 8,121 | ZIP |
| Vinblastine | 5,148 | 14,378 | ZIP |
| Upadacitinib | 5,119 | 8,189 | ZIP |

| | | | |
|---|---|---|---|
| Orteronel | 5,113 | 10,149 | ZIP |
| Canertinib | 5,098 | 8,715 | ZIP |
| XAV-939 | 5,082 | 8,366 | ZIP |
| BAY 87-2243 | 5,072 | 10,48 | ZIP |
| UNC0642 | 5,066 | 10,492 | ZIP |
| Belinostat | 5,039 | 13,404 | ZIP |
| Onalespib | 5,012 | 8,39 | ZIP |
| Metformin | 4,999 | 7,471 | ZIP |
| AZD4547 | 4,986 | 10,419 | ZIP |
| UM729 | 4,977 | 10,726 | ZIP |
| Mocetinostat | 4,971 | 8,915 | ZIP |
| Tacedinaline | 4,965 | 7,771 | ZIP |
| Ralimetinib | 4,963 | 9,98 | ZIP |
| Lomeguatrib | 4,933 | 9,528 | ZIP |
| Pevonedistat | 4,93 | 14,179 | ZIP |
| Entinostat | 4,902 | 10,26 | ZIP |
| Infigratinib | 4,893 | 10,182 | ZIP |
| Mitomycin C | 4,866 | 8,793 | ZIP |
| Miltefosine | 4,857 | 9,136 | ZIP |
| GSK2879552 | 4,812 | 7,471 | ZIP |
| ODM-201 | 4,806 | 9,368 | ZIP |
| Quisinostat | 4,806 | 8,876 | ZIP |
| Fedratinib | 4,78 | 8,93 | ZIP |
| GSK2256098 | 4,779 | 8,577 | ZIP |
| Ganetespib | 4,747 | 8,116 | ZIP |
| A-366 | 4,745 | 8,403 | ZIP |
| PF06650833 | 4,739 | 9,132 | ZIP |
| VS-4718 | 4,729 | 8,908 | ZIP |
| Erastin | 4,703 | 11,126 | ZIP |
| Pracinostat | 4,7 | 10,731 | ZIP |
| MST-312 | 4,681 | 13,087 | ZIP |
| AMG-925 | 4,648 | 7,434 | ZIP |
| TEW-7197 | 4,646 | 13,91 | ZIP |
| Amcasertib | 4,622 | 11,533 | ZIP |
| Verdinexor | 4,596 | 8,091 | ZIP |
| 1-methyl-D-tryptophan | 4,571 | 10,415 | ZIP |
| ML323 | 4,56 | 8,887 | ZIP |
| Romidepsin | 4,553 | 10,705 | ZIP |
| Plicamycin | 4,539 | 10,586 | ZIP |
| AR-42 | 4,538 | 11,285 | ZIP |
| ASP3026 | 4,515 | 8,547 | ZIP |
| AZ191 | 4,515 | 7,778 | ZIP |

| | | | |
|---|---|---|---|
| MK-8776 | 4,494 | 11,468 | ZIP |
| Ponatinib | 4,493 | 10,359 | ZIP |
| AZD7762 | 4,481 | 6,957 | ZIP |
| Volasertib | 4,459 | 9,353 | ZIP |
| Rociletinib | 4,446 | 7,453 | ZIP |
| TG100-115 | 4,431 | 9,245 | ZIP |
| Dacomitinib | 4,415 | 8,76 | ZIP |
| Toremifene | 4,406 | 5,831 | ZIP |
| PCI-34051 | 4,402 | 8,429 | ZIP |
| Olmutinib | 4,397 | 7,76 | ZIP |
| AT7519 | 4,395 | 12,406 | ZIP |
| AZD6738 | 4,376 | 8,155 | ZIP |
| I-BET151 | 4,375 | 10,866 | ZIP |
| Cladribine | 4,369 | 10,059 | ZIP |
| 8-chloro-adenosine | 4,367 | 7,785 | ZIP |
| Apalutamide | 4,36 | 7,944 | ZIP |
| CCT196969 | 4,347 | 8,218 | ZIP |
| Neflamapimod | 4,345 | 6,095 | ZIP |
| BMS-911543 | 4,34 | 8,445 | ZIP |
| Pilocarpine | 4,333 | 10,008 | ZIP |
| NVP-RAF265 | 4,308 | 7,766 | ZIP |
| PAC-1 | 4,29 | 10,871 | ZIP |
| AVN944 | 4,286 | 7,221 | ZIP |
| TIC10 | 4,273 | 8,733 | ZIP |
| Tideglusib | 4,255 | 9,147 | ZIP |
| Tivantinib | 4,243 | 9,164 | ZIP |
| AT-406 | 4,22 | 10,215 | ZIP |
| GSK-2334470 | 4,216 | 10,024 | ZIP |
| Tamoxifen | 4,201 | 10,326 | ZIP |
| Tozasertib | 4,195 | 8,522 | ZIP |
| Pinometostat | 4,183 | 8,107 | ZIP |
| Rocilinostat | 4,167 | 10,607 | ZIP |
| LY-2584702 | 4,133 | 8,014 | ZIP |
| Pacritinib | 4,109 | 6,815 | ZIP |
| AZD-6482 | 4,101 | 8,513 | ZIP |
| Losmapimod | 4,099 | 10,834 | ZIP |
| Idelalisib | 4,075 | 9,024 | ZIP |
| PF-3845 | 4,045 | 8,336 | ZIP |
| DEL-22379 | 4,039 | 8,05 | ZIP |
| 8-amino-adenosine | 4,017 | 10,532 | ZIP |
| Bosutinib | 3,994 | 6,221 | ZIP |
| Cediranib | 3,99 | 7,968 | ZIP |

| | | | |
|---|---|---|---|
| Enzalutamide | 3,978 | 6,45 | ZIP |
| Tucidinostat | 3,972 | 11,825 | ZIP |
| BI 2536 | 3,956 | 6,174 | ZIP |
| Vandetanib | 3,952 | 7,358 | ZIP |
| Ensartinib | 3,95 | 8,359 | ZIP |
| AZD1208 | 3,937 | 6,356 | ZIP |
| UNC0638 | 3,908 | 7,638 | ZIP |
| OTS167 | 3,907 | 7,252 | ZIP |
| Spebrutinib | 3,902 | 8,201 | ZIP |
| Nintedanib | 3,887 | 6,685 | ZIP |
| Cabozantinib | 3,868 | 6,067 | ZIP |
| Triapine | 3,824 | 8,56 | ZIP |
| Roxadustat | 3,808 | 6,697 | ZIP |
| Osimertinib | 3,803 | 12,13 | ZIP |
| Rucaparib | 3,779 | 8,203 | ZIP |
| Erdafitinib | 3,755 | 7,49 | ZIP |
| Talazoparib | 3,742 | 4,618 | ZIP |
| Oprozomib | 3,736 | 6,474 | ZIP |
| IOX-2 | 3,722 | 7,321 | ZIP |
| UNC2881 | 3,709 | 9,078 | ZIP |
| Saracatinib | 3,695 | 6,457 | ZIP |
| Sunitinib | 3,68 | 8,77 | ZIP |
| Erlotinib | 3,677 | 9,563 | ZIP |
| BRD7116 | 3,673 | 8,946 | ZIP |
| Buparlisib | 3,667 | 6,136 | ZIP |
| NVP-AEW541 | 3,637 | 8,556 | ZIP |
| Sapitinib | 3,619 | 8,01 | ZIP |
| Givinostat | 3,61 | 6,629 | ZIP |
| LY3009120 | 3,587 | 7,6 | ZIP |
| Auranofin | 3,583 | 7,79 | ZIP |
| TGX-221 | 3,575 | 8,374 | ZIP |
| Epacadostat | 3,555 | 7,606 | ZIP |
| KU-60019 | 3,553 | 7,999 | ZIP |
| AMG-337 | 3,541 | 7,593 | ZIP |
| KD025 | 3,539 | 11,266 | ZIP |
| SH-4-54 | 3,515 | 7,694 | ZIP |
| Seliciclib | 3,5 | 9,54 | ZIP |
| Disulfiram(+CuCl2) | 3,468 | 8,475 | ZIP |
| SCH772984 | 3,46 | 8,777 | ZIP |
| Momelotinib | 3,46 | 8,519 | ZIP |
| Everolimus | 3,434 | 4,248 | ZIP |
| Varespladib | 3,416 | 7,818 | ZIP |

| | | | |
|---|---|---|---|
| Darapladib | 3,409 | 7,228 | ZIP |
| Alisertib | 3,4 | 8,141 | ZIP |
| Vinflunine | 3,399 | 8,724 | ZIP |
| PF-4800567 | 3,395 | 7,149 | ZIP |
| JQ1 | 3,373 | 7,198 | ZIP |
| GDC-0623 | 3,371 | 7,752 | ZIP |
| EPZ031686 | 3,345 | 7,864 | ZIP |
| Clomifene | 3,338 | 6,055 | ZIP |
| Decernotinib | 3,329 | 9,014 | ZIP |
| TAK-530 | 3,325 | 10,219 | ZIP |
| Idarubicin | 3,285 | 5,729 | ZIP |
| Afatinib | 3,267 | 8,953 | ZIP |
| Tesevatinib | 3,261 | 4,935 | ZIP |
| Lenvatinib | 3,238 | 8,3 | ZIP |
| Sapanisertib | 3,234 | 8,369 | ZIP |
| Copanlisib | 3,227 | 5,773 | ZIP |
| Quizartinib | 3,226 | 5,096 | ZIP |
| Clofarabine | 3,22 | 9,023 | ZIP |
| Topotecan | 3,197 | 6,919 | ZIP |
| Poziotinib | 3,176 | 6,524 | ZIP |
| BGB-283 | 3,171 | 5,93 | ZIP |
| Napabucasin | 3,152 | 7,117 | ZIP |
| Resatorvid | 3,133 | 7,411 | ZIP |
| Olaparib | 3,128 | 3,988 | ZIP |
| Galiellalactone | 3,108 | 9,843 | ZIP |
| Floxuridine | 3,099 | 6,823 | ZIP |
| Sabutoclax | 3,094 | 5,307 | ZIP |
| Vesatolimod | 3,077 | 6,717 | ZIP |
| Cytarabine/Idarubicin | 3,075 | 8,617 | ZIP |
| Uprosertib | 3,075 | 5,599 | ZIP |
| Glasdegib | 3,06 | 10,91 | ZIP |
| Aldoxorubicin | 3,054 | 6,162 | ZIP |
| Omaveloxolone | 3,043 | 6,152 | ZIP |
| GDC-0919 | 3,04 | 9,185 | ZIP |
| Atorvastatin | 3 | 6,771 | ZIP |
| Selonsertib | 2,985 | 9,191 | ZIP |
| WEHI-539 | 2,956 | 8,087 | ZIP |
| Ceritinib | 2,943 | 7,646 | ZIP |
| Entrectinib | 2,939 | 5,592 | ZIP |
| Vistusertib | 2,938 | 3,544 | ZIP |
| Anastrozole | 2,889 | 10,401 | ZIP |
| BIIB021 | 2,867 | 6,019 | ZIP |

| | | | |
|---|---|---|---|
| Dasatinib | 2,846 | 5,154 | ZIP |
| AT9283 | 2,809 | 5,981 | ZIP |
| Axitinib | 2,762 | 9,631 | ZIP |
| Crenolanib | 2,758 | 5,389 | ZIP |
| BMS863233 | 2,755 | 7,814 | ZIP |
| Gandotinib | 2,744 | 7,076 | ZIP |
| Cobimetinib | 2,741 | 5,017 | ZIP |
| Vorinostat | 2,735 | 7,265 | ZIP |
| AZD-5363 | 2,733 | 4,577 | ZIP |
| Ruxolitinib | 2,72 | 4,535 | ZIP |
| Acitretin | 2,705 | 4,621 | ZIP |
| Lasofoxifene | 2,658 | 12,078 | ZIP |
| Lenalidomide | 2,65 | 8,927 | ZIP |
| A-1155463 | 2,629 | 5,496 | ZIP |
| Gemcitabine | 2,619 | 13,861 | ZIP |
| Pentostatin | 2,616 | 5,458 | ZIP |
| ML390 | 2,612 | 5,399 | ZIP |
| Cilengitide | 2,594 | 6,814 | ZIP |
| Cytarabine | 2,572 | 8,604 | ZIP |
| Peficitinb | 2,565 | 7,608 | ZIP |
| Tubastatin A | 2,564 | 7,541 | ZIP |
| Gefitinib | 2,553 | 8,071 | ZIP |
| Encorafenib | 2,551 | 6,951 | ZIP |
| LY-2874455 | 2,535 | 5,896 | ZIP |
| VGX-1027 | 2,529 | 7,867 | ZIP |
| NVP-SHP099 | 2,492 | 7,473 | ZIP |
| Sepantronium bromide | 2,484 | 5,315 | ZIP |
| RO5126766 | 2,469 | 6,949 | ZIP |
| URB597 | 2,467 | 6,185 | ZIP |
| Ribociclib | 2,466 | 3,99 | ZIP |
| Golvatinib | 2,457 | 5,072 | ZIP |
| Glesatinib | 2,456 | 6,561 | ZIP |
| Carboplatin | 2,425 | 5,532 | ZIP |
| Valrubicin | 2,411 | 5,785 | ZIP |
| Ixabepilone | 2,392 | 5,423 | ZIP |
| Merestinib | 2,39 | 6,188 | ZIP |
| Varlitinib | 2,388 | 7,42 | ZIP |
| PF-670462 | 2,378 | 7,477 | ZIP |
| TAK-285 | 2,37 | 5,743 | ZIP |
| E7820 | 2,322 | 7,374 | ZIP |
| Brivanib | 2,314 | 5,475 | ZIP |
| RSL3 | 2,282 | 11,905 | ZIP |

| | | | |
|---|---|---|---|
| Amuvatinib | 2,269 | 7,183 | ZIP |
| Carfilzomib | 2,258 | 5,313 | ZIP |
| Ulixertinib | 2,256 | 4,198 | ZIP |
| Bicalutamide | 2,228 | 5,101 | ZIP |
| ONX-0914 | 2,227 | 5,168 | ZIP |
| AZ 3146 | 2,187 | 6,427 | ZIP |
| Oxaliplatin | 2,155 | 8,638 | ZIP |
| Doramapimod | 2,146 | 7,19 | ZIP |
| Digoxin | 2,125 | 4,578 | ZIP |
| Daporinad | 2,111 | 4,599 | ZIP |
| MK-2206 | 2,107 | 5,389 | ZIP |
| AMG319 | 2,105 | 5,963 | ZIP |
| Gedatolisib | 2,08 | 6,378 | ZIP |
| Alectinib | 2,036 | 6,234 | ZIP |
| Nelarabine | 2,035 | 3,46 | ZIP |
| AZD-1080 | 2,005 | 5,837 | ZIP |
| A-1210477 | 1,986 | 6,552 | ZIP |
| AZD1775 | 1,982 | 4,588 | ZIP |
| MK-8745 | 1,958 | 5,206 | ZIP |
| Marimastat | 1,94 | 5,105 | ZIP |
| Tazemetostat | 1,919 | 4,641 | ZIP |
| Vemurafenib | 1,89 | 6,591 | ZIP |
| Fingolimod | 1,86 | 3,369 | ZIP |
| Mepacrine | 1,841 | 5,282 | ZIP |
| Ipatasertib | 1,825 | 4,44 | ZIP |
| Lonafarnib | 1,763 | 6,395 | ZIP |
| Capmatinib | 1,686 | 7,283 | ZIP |
| Lovastatin | 1,673 | 9,906 | ZIP |
| Daunorubicin | 1,665 | 5,985 | ZIP |
| Exemestane | 1,639 | 3,912 | ZIP |
| Motolimod | 1,604 | 8,584 | ZIP |
| Selumetinib | 1,594 | 4,578 | ZIP |
| AZD3759 | 1,583 | 9,105 | ZIP |
| Palbociclib | 1,577 | 3,803 | ZIP |
| Palomid-529 | 1,574 | 8,609 | ZIP |
| Dactolisib | 1,554 | 3,25 | ZIP |
| Capecitabine | 1,548 | 3,766 | ZIP |
| AZD1152-HQPA | 1,542 | 3,562 | ZIP |
| Dovitinib | 1,514 | 3,626 | ZIP |
| CEP-37440 | 1,511 | 3,368 | ZIP |
| GNE-0877 | 1,501 | 5,861 | ZIP |
| Brigatinib | 1,489 | 4,558 | ZIP |

| | | | |
|---|---|---|---|
| APR-246 | 1,441 | 2,407 | ZIP |
| CC-223 | 1,389 | 5,919 | ZIP |
| Thioguanine | 1,389 | 3,865 | ZIP |
| Masitinib | 1,376 | 2,553 | ZIP |
| Bortezomib | 1,364 | 3,372 | ZIP |
| Filgotinib | 1,336 | 4,254 | ZIP |
| Nilutamide | 1,328 | 3,41 | ZIP |
| VER 155008 | 1,238 | 5,04 | ZIP |
| EPZ-5687 | 1,223 | 5,997 | ZIP |
| Methotrexate | 1,199 | 2,933 | ZIP |
| Crizotinib | 1,187 | 4,737 | ZIP |
| SGI-1776 | 1,185 | 5,141 | ZIP |
| Tandutinib | 1,18 | 5,985 | ZIP |
| GDC-0853 | 1,16 | 4,347 | ZIP |
| Epirubicin | 1,137 | 4,604 | ZIP |
| Simvastatin | 1,098 | 3,175 | ZIP |
| Imatinib | 1,086 | 7,432 | ZIP |
| Danusertib | 1,069 | 6,758 | ZIP |
| Galunisertib | 1,042 | 2,828 | ZIP |
| Venetoclax | 0,972 | 6,985 | ZIP |
| Celecoxib | 0,954 | 6,841 | ZIP |
| GNE-7915 | 0,889 | 4,851 | ZIP |
| TGR-1202 | 0,863 | 3,297 | ZIP |
| A-1331852 | 0,861 | 3,517 | ZIP |
| Pazopanib | 0,783 | 3,635 | ZIP |
| Entospletinib | 0,759 | 7,817 | ZIP |
| Ixazomib | 0,749 | 2,293 | ZIP |
| Pirfenidone | 0,747 | 2,142 | ZIP |
| JPH203 | 0,74 | 7,056 | ZIP |
| Necrostatin 2 | 0,732 | 4,888 | ZIP |
| ZSTK474 | 0,687 | 6,316 | ZIP |
| Midostaurin | 0,669 | 4,107 | ZIP |
| IOX-1 | 0,66 | 3,83 | ZIP |
| AT 101 | 0,655 | 7,567 | ZIP |
| LY3023414 | 0,643 | 2,731 | ZIP |
| PH-797804 | 0,628 | 5,508 | ZIP |
| GSK-J4 | 0,552 | 6,503 | ZIP |
| TH588 | 0,448 | 7,388 | ZIP |
| Letrozole | 0,447 | 2,795 | ZIP |
| Tarenflurbil | 0,438 | 5,883 | ZIP |
| Bimatoprost | 0,429 | 5,547 | ZIP |
| Panobinostat | 0,426 | 15,362 | ZIP |

| | | | |
|---|---|---|---|
| Enzastaurin | 0,402 | 2,988 | ZIP |
| OSU-03012 | 0,36 | 5,261 | ZIP |
| Linsitinib | 0,349 | 3,446 | ZIP |
| PD0325901 | 0,297 | 3,547 | ZIP |
| Binimetinib | 0,288 | 1,635 | ZIP |
| Ravoxertinib | 0,282 | 3,415 | ZIP |
| Tasquinimod | 0,038 | 4,396 | ZIP |
| Motesanib | -0,01 | 1,46 | ZIP |
| AZD7545 | -0,053 | 6,804 | ZIP |
| Arsenic(III) oxide | -0,099 | 8,602 | ZIP |
| Pomalidomide | -0,1 | 1,859 | ZIP |
| Teniposide | -0,104 | 7,065 | ZIP |
| MK-0752 | -0,112 | 5,65 | ZIP |
| Taladegib | -0,152 | 4,413 | ZIP |
| Valproic acid | -0,23 | 3,027 | ZIP |
| Regorafenib | -0,267 | 2,371 | ZIP |
| Ruboxistaurin | -0,273 | 3,447 | ZIP |
| Lapatinib | -0,287 | 5,713 | ZIP |
| GSK2830371 | -0,313 | 4,891 | ZIP |
| Chloroquine | -0,38 | 4,926 | ZIP |
| Azacitidine | -0,442 | 5,807 | ZIP |
| Temozolomide | -0,445 | 5,059 | ZIP |
| Bentamapimod | -0,456 | 0,405 | ZIP |
| CC122 | -0,585 | 4,722 | ZIP |
| Mercaptopurine | -0,65 | 1,689 | ZIP |
| Bafetinib | -0,666 | 1,645 | ZIP |
| Tirabrutinib | -0,73 | 3,007 | ZIP |
| Sonolisib | -0,757 | 3,584 | ZIP |
| Hydroxyurea | -0,792 | 1,786 | ZIP |
| Plerixafor | -0,797 | 7,288 | ZIP |
| Raltitrexed | -0,842 | 3,564 | ZIP |
| EPZ015666 | -0,87 | 3,327 | ZIP |
| Imiquimod | -1,05 | 0,998 | ZIP |
| Dabrafenib | -1,061 | 5,552 | ZIP |
| Megestrol acetate | -1,064 | 1,712 | ZIP |
| VLX1570 | -1,116 | 4,089 | ZIP |
| Radotinib | -1,316 | 0,93 | ZIP |
| Finasteride | -1,382 | 5,665 | ZIP |
| Tretinoin | -1,407 | 0,806 | ZIP |
| Goserelin | -1,46 | 3,204 | ZIP |
| NVP-LGK974 | -1,55 | 8,291 | ZIP |
| Fluorouracil | -1,642 | 1,598 | ZIP |

| | | | |
|---|---|---|---|
| VE-821 | -1,652 | 2,274 | ZIP |
| Apatinib | -1,751 | 9,396 | ZIP |
| Navitoclax | -1,754 | 2,45 | ZIP |
| Tacrolimus | -1,812 | 0,265 | ZIP |
| Raloxifene | -1,882 | 3,164 | ZIP |
| Trametinib | -1,901 | 5,108 | ZIP |
| Acalabrutinib | -1,917 | 0,642 | ZIP |
| Hydroxyfasudil | -2,171 | 7,765 | ZIP |
| Sonidegib | -2,273 | 0,011 | ZIP |
| Aminoglutethimide | -2,299 | -0,088 | ZIP |
| Sotrastaurin | -2,442 | 1,708 | ZIP |
| Bryostatin 1 | -2,476 | 3,426 | ZIP |
| Vatalanib | -2,491 | 2,655 | ZIP |
| StemRegenin 1 | -2,616 | 4,527 | ZIP |
| Abiraterone | -2,753 | 2,342 | ZIP |
| Asciminib | -2,782 | 2,197 | ZIP |
| Mubritinib | -2,805 | 2,81 | ZIP |
| Fulvestrant | -2,873 | 4,923 | ZIP |
| Telatinib | -2,92 | -0,283 | ZIP |
| Anagrelide | -2,934 | 4,468 | ZIP |
| UNC1215 | -3,178 | 0,07 | ZIP |
| Enasidenib | -3,2 | -0,39 | ZIP |
| Perifosine | -3,289 | -1,747 | ZIP |
| Decitabine | -3,297 | 1,555 | ZIP |
| PS-1145 | -3,374 | -1,17 | ZIP |
| Pexidartinib | -3,456 | 3,048 | ZIP |
| Bexarotene | -3,556 | -0,164 | ZIP |
| Larotrectinib | -3,583 | 0,02 | ZIP |
| CEP-32496 | -4,265 | 0,897 | ZIP |
| Trifluridine | -4,297 | 6,441 | ZIP |
| Linifanib | -4,518 | -0,769 | ZIP |
| Nilotinib | -4,761 | -2,201 | ZIP |
| Vismodegib | -5,597 | 1,094 | ZIP |
| Methylprednisolone | -5,631 | -1,07 | ZIP |
| Salinomycin | -5,666 | 14,631 | ZIP |
| Ibrutinib | -6,309 | -1,523 | ZIP |
| Thalidomide | -6,542 | -3,751 | ZIP |
| Flutamide | -6,589 | 2,773 | ZIP |
| Dexamethasone | -8,016 | -3,553 | ZIP |
| Allopurinol | -8,018 | -4,049 | ZIP |
| AZD3965 | -8,464 | -1,937 | ZIP |
| Senexin B | -9,096 | -4,201 | ZIP |

| | | | |
|---|---|---|---|
| Sorafenib | -9,47 | -4,504 | ZIP |
| Prednisolone | -10,938 | -4,015 | ZIP |
| Mitotane | -11,142 | -5,904 | ZIP |

Table S3. **The developmental status of Bcl-2 inhibitors in combinations with radiotherapy.**

| **Bcl2i** | **Reference** | **Developmental stage** | **Condition** |
|---|---|---|---|
| ABT-263 | PMID:30614795 | SCLC cell lines | SCLC |
| ABT-737 | PMID:31579427 | uterine cervical cancer cells | uterine cervical cancer |
| ABT-737 | PMID:26934442 | HNSCC cell lines | HNSCC |
| ABT-737 | PMID:25409124 | breast cancer cell lines | breast cancer |
| ABT-737 | PMID:23285061 | cervical cancer HeLa cells | cervical cancer |
| ABT-737 | PMID:23259599 | breast cancer cells | breast cancer |
| ABT-737 | PMID:23259599 | breast cancer cells | breast cancer |
| ABT-737 | PMID:22002102 | glioblastoma cells | glioblastoma |
| Gossypol | PMID:19852810 | human leukemic cells | leukemia |
| Gossypol | PMID:17521756 | tumour cell lines | tumors |
| Gossypol | PMID:15713891 | human prostate cancer cells | prostate cancer |
| Gossypol | PMID:26223311 | HNSCC cell lines | HNSCC |
| Gossypol | PMID:24968413 | radioresistant malignant glioma | Malignant gliomas |
| Gossypol | PMID:21229643 | prostate cancer cells | prostate cancer |
| Gossypol | PMID:21319440 | human prostate cancer cells | prostate cancer |
| Gossypol | PMID:20354451 | lung cancer cells | lung cancer |
| Gossypol | NCT00390403 | phase I trial | Newly Diagnosed Glioblastoma Multiforme |
| Obatoclax | PMID:25568669 | glioblastoma stem-like cells | glioblastoma |
| TW-37 | PMID:20675079 | tumor angiogenesis in vivo | HNSCC |
| HA14-1 | PMID:18774194 | cervical cancer cells | cervical cancer |
| Gamboic acid | PMID:26357974 | nasopharyngeal carcinoma cells | NPC |
| Gamboic acid | PMID:26318432 | esophageal cancer cells | Esophageal cancer |
| BH3I-1 | PMID:15909480 | NSCC cells | NSCC |
| ABT-199 | PMID:28566329 | xenograft models of lymphomas | B cell Lymphomas |
